# Two-component systems regulate swarming in *Pseudomonas aeruginosa* PA14

**DOI:** 10.1101/445015

**Authors:** Shubham Joge, Ameen M. Kollaran, Harshitha S. Kotian, Divakar Badal, Deep Prakash, Ayushi Mishra, Manoj Varma, Varsha Singh

## Abstract

Swarming in *Pseudomonas aeruginosa* is a quorum-dependant motility over semi-solid surfaces. On soft agar, *P. aeruginosa* exhibits a dendritic swarm pattern, with multiple levels of branching. Swarm patterns vary considerably depending upon the experimental design. In the present study, we show that the swarm pattern is plastic and media dependent. We define several quantifiable, macroscale features of the swarm to study the plasticity observed across media. Further, through a targeted screen of 113 genes encoding two-component system (TCS) components, we show that 44 TCS genes regulate PA14 swarming in a contextual fashion. However, only four TCS genes are essential for swarming. Many swarming-defective TCS mutants are highly efficient in biofilm formation indicating an antagonistic relationship between swarming and biofilm states in *P. aeruginosa*.

## INTRODUCTION

Swarming is a rapid bulk translocation behaviour observed in many bacterial species, typically over semi-solid agar surfaces (Harshey & Matsuyama, 1994; Henrichsen, 1972; Kearns, 2010). In several instances, bacterial swarm populations exhibit a characteristic macroscopic swarm patterns, that are easily recognizable (Kearns, 2010). The swarm patterns are often species-specific and can be classified as circular, vortex, featureless, branched or composed of concentric whorls (Kearns 2010). Other than a few exceptions like *Myxococcus xanthus*, most swarming bacteria are bacilli which rely on flagella for swarming (Kaiser & Warrick, 2011; Kearns, 2010).

*Pseudomonas aeruginosa* is a Gram-negative, flagellated bacillus which displays branched or dendritic swarm pattern on soft agar. Other than flagella, it requires rhamnolipid surfactants for swarming (Kohler et al., 2000; Overhage, Bains, Brazas, & Hancock, 2008). Rhamnolipids, a class of glycolipid biosurfactants, are a critical factor in *P. aeruginosa* swarming, including tendril avoidance in the swarm (Caiazza, Shanks, & Toole, 2005; Morris et al., 2011, Xavier, Kim, & Foster, 2011). Rhamnolipid production in *P. aeruginosa* can be regulated by nitrogen source in M8 minimal medium (Kohler et al., 2000) and by carbon sources (Shrout et al., 2006). Rhamnolipid production can also be induced in *P. aeruginosa* PAO1 by phosphate limitation in BM2 minimal medium (Bains M, Fernández L, 2012). Both carbon or nitrogen sources are also known to affect the ability of *P. aeruginosa* to swarm (Kohler et al., 2000; Shrout et al., 2006). Most *P. aeruginosa* swarming are primarily reported under minimal media conditions (M8, M9 and BM2) or in some instance, under complex media formulations such as nutrient broth, brain heart infusion (BHI) or fastidious anaerobe broth (FAB) (Baker et al., 2016; Kohler et al., 2000; Morales-Soto et al., 2015; Overhage et al., 2008; Rashid & Kornberg, 2000; Tremblay & Déziel, 2008). These media vary in concentration of various macro as well as of micronutrients. However, despite a gross conservation in the dendritic swarm pattern on different media, the *P. aeruginosa* swarms appear distinct. These observations raised the question of whether nutritional components of the growth medium affect *P. aeruginosa* swarm ability and pattern.

Transcriptome analysis of *P. aeruginosa* swarming has been shown to result in upregulation of virulence factors on BM2 medium (Overhage et al., 2008) whereas another study showed dysregulation of several components of the translation machinery, energy metabolism and nutrient transporters on M9 medium (Tremblay & Déziel, 2010). Such a notable divergence between these two studies might have resulted from the use of two different media (BM2 and M9) which differ in nutritional composition (Tremblay & Déziel, 2010). There also exists at least one instance where a *P. aeruginosa* mutant displays contrasting swarming phenotypes. A pili mutant, *pilA*, of *P. aeruginosa* is described as a non-swarmer on M8 agar (Kohler et al., 2000), reported a swarmer on nutrient broth agar (Rashid & Kornberg, 2000) and hyper-swarmer on FAB agar (Shrout, 2006). These observations also suggested the possible impact of nutrient components on the *P. aeruginosa* swarming, and upon the requirement of genetic regulators. An interesting question is how *P. aeruginosa* cells sense changes in nutrition that impacts swarm phenotype. However, a comprehensive analysis of the possible impact of various nutrients or the importance of bacterial nutrient sensors in *P. aeruginosa* swarming is yet to be carried out. In the present study, we analysed swarming behaviour of *P. aeruginosa* strain PA14 across six different nutrient agars and define swarm features that can be quantified easily. We show that swarm patterns vary considerably across media with reproducible, medium-specific features. We also show that forty-four genes encoding two-component systems (TCSs) including several poorly characterised or unstudied sensor kinases and response regulators are required for *P. aeruginosa* swarming. Among these, four TCS genes are essential for swarming on all media, while the remaining have context specific function. We also find that several positive regulators of swarming have an opposite effect on biofilm formation.

## RESULTS

### Phenotypic plasticity *in P. aeruginosa* swarming is nutrition dependent

To understand whether nutrition has an impact on swarming, we analysed *P. aeruginosa* swarming on six different media – Luria Bertani (LB), Brain Heart Infusion (BHI), M8, M9, PGM and BM2 – the nutrient formulations often described for *P. aeruginosa* growth or swarming studies (Kohler et al., 2000; Morris et al., 2011; Overhage et al., 2008, 2007; Yeung et al., 2009) (Table S1). Peptone growth media (PGM), also called slow killing medium, is used for *P. aeruginosa* growth in *Caenorhabditis elegans* infection studies (Singh & Aballay, 2006; Sun et al., 2011; Tan, Mahajan-Miklos, & Ausubel, 1999). As shown in Figure 1A-F, we found that four media supported dendritic swarm pattern for *P. aeruginosa* PA14. However, BHI and LB media did not support dendrite formation, a characteristic of *P. aeruginosa* swarming (Figure 1A, B). LB and BHI agar also poorly supported isometric swarm expansion pattern exhibited by other species such as *Escherichia coli, Salmonella typhimurium* or *Bacillus subtilis* (Harshey & Matsuyama, 1994; Kearns & Losick, 2003; Patrick & Kearns, 2009). Thus, three minimal media (M8, M9, and BM2) and one undefined medium (PGM) supported swarming with distinct dendrites. All four media also supported multiple (1-3) levels of branching (Figure 1, C-F) but had medium specific or plastic patterns.

**Figure 1.**
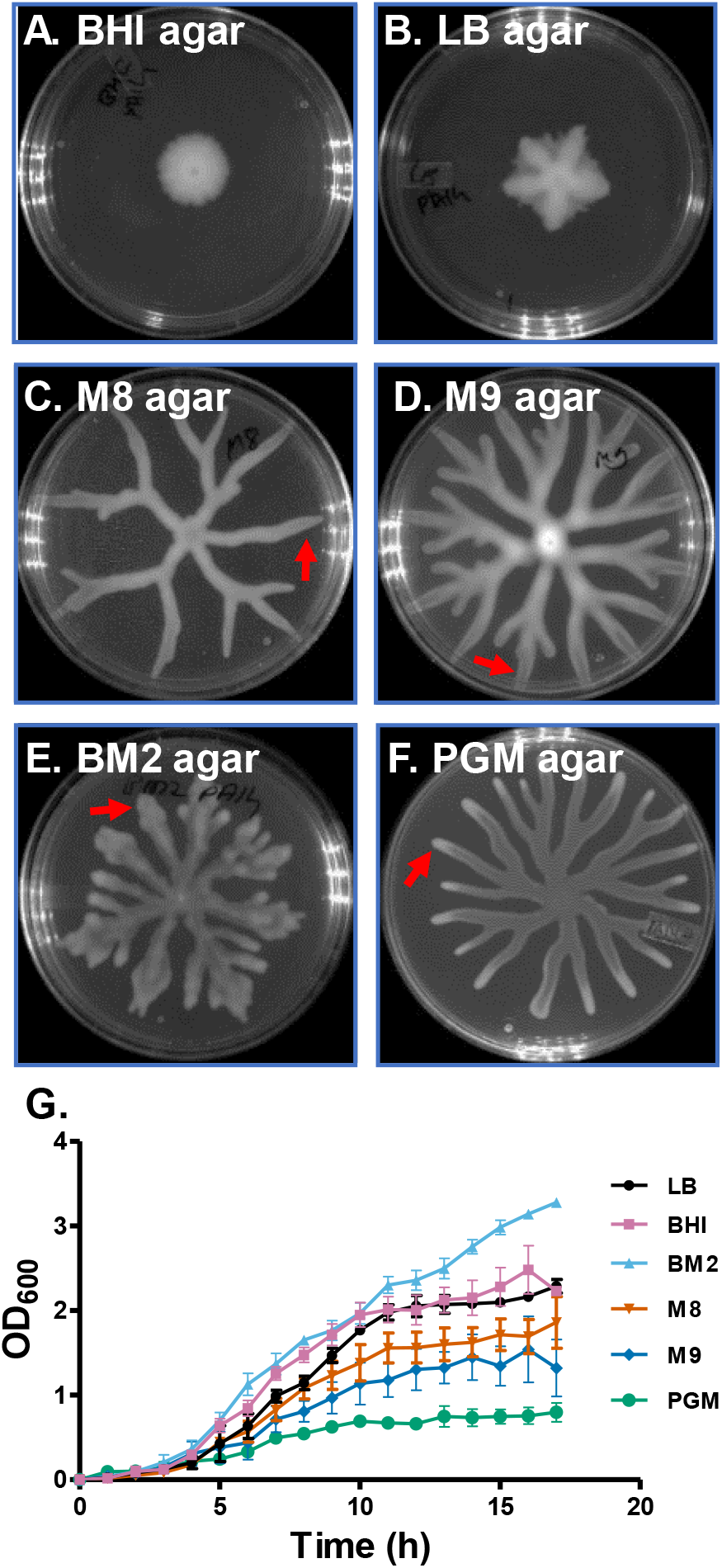
*P. aeruginosa* PA14 swarm pattern on different swarming agar. ***P** aeruginosa* swarm obtained at 37°C for 24hours on (**A**) brain heart infusion (BHI), (**B**) Luria Bertani (LB), (**C**) M8, (**D**) M9, (**E**) BM2 and, (**F**) Peptone growth media (PGM) with 0.6% agar. (**G**) PA14 WT planktonic growth kinetics on various media is shown. Dendrites are indicated with arrow in C-F.

Further, we analysed the planktonic growth kinetics of PA14 in all six-media, mentioned above. Both LB and BHI broth supported better growth of PA14 (Figure 1G) compared to M8, M9, and PGM broth. The BM2 broth was also able to support good planktonic growth like LB and BHI broth. M8 and M9 supported moderate growth while PGM supported poor growth for PA14. Hence, except for BM2 medium, our data suggest an inverse relationship between the planktonic growth and the propensity for swarming in *P. aeruginosa* PA14. Taken together, analysis of swarm agar and broth phase growth of PA14 in three undefined media and three minimal media suggests that poor media, probably lacking specific nutrients, promote swarming. This data also provides evidence for nutrition dependent plasticity in *P. aeruginosa* swarm pattern.

### Macroscale features define the plasticity of *P. aeruginosa* swarm

To characterise the plasticity in PA14 swarm patterns across media, we set out to define features of the swarm that could be quantified. We found that the conventional approach of comparing a single feature such as the swarm diameter or bacterial cell number (Overhage et al., 2007; Xavier, Kim, & Foster, 2011; Yeung et al., 2009) was not suitable to differentiate dendritic patterns observed on four different media in our study (Figure 1C-F). By analysing several swarm images for each of the media, we defined eleven measurable features of *P. aeruginosa* swarm illustrated in Figure 2A-C. These included branch angle, branch width, number of growing tips, area of the swarm, swarm perimeter, normalized area etc (Figure 2A, B, see methods). Swarm-lag (the time taken to initiate branching from the time of spotting) is the shortest on M9 followed by M8, BM2 and PGM agar (Video S1 to S6). This is represented in Figure 2C as circularity versus time. Drop in circularity below the value of 1.0 marks the end of the swarm lag and initiation of branching. We find that branching starts first in M9, followed by M8 and then on BM2 agar. The swarm lag is longest on PGM agar (Figure 2C). Our observations suggest that medium influences regulatory program that dictates the set time to initiate branching.

**Figure 2.**
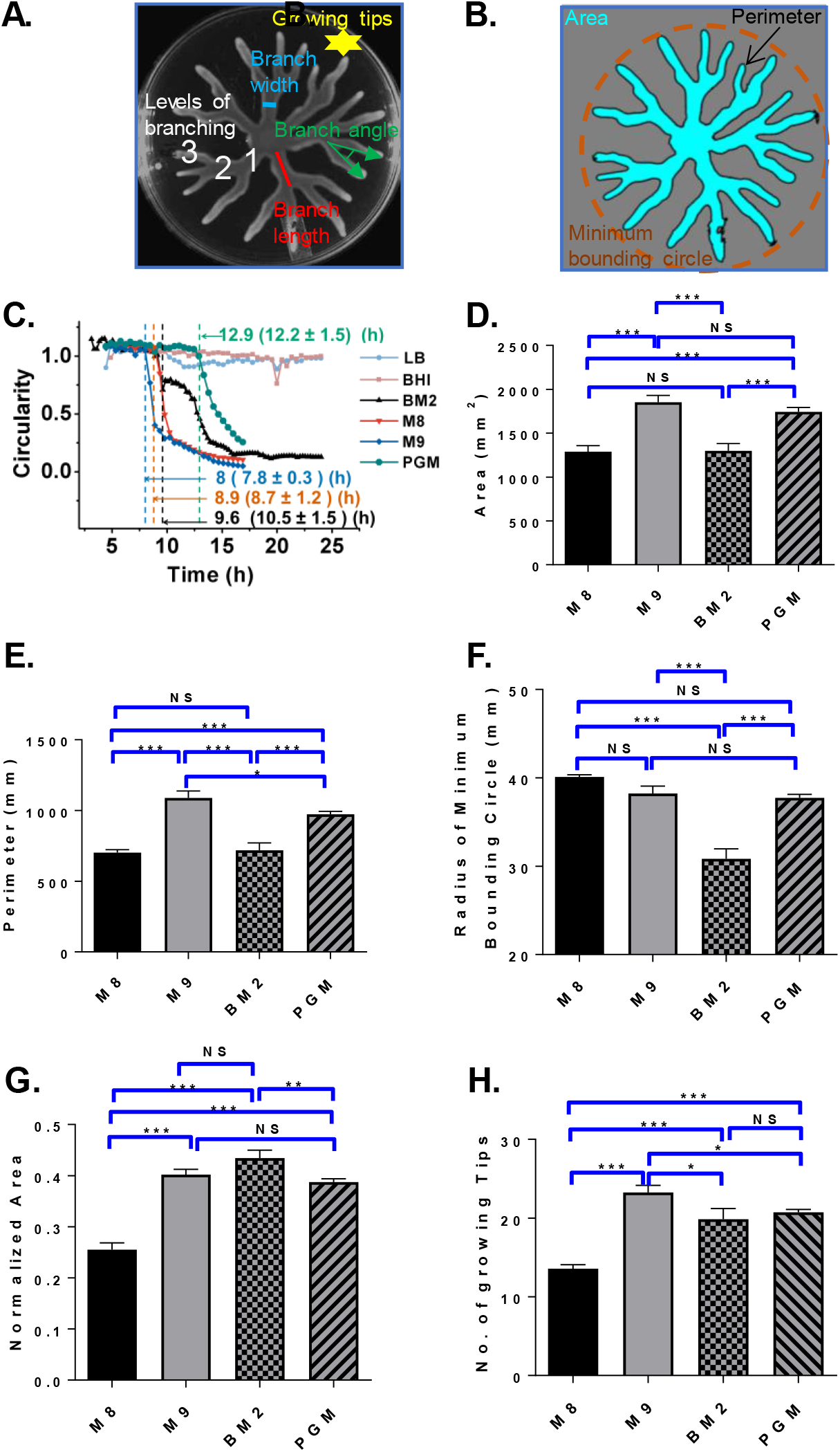
Macroscale features of *P. aeruginosa* swarm. (**A-B**) Quantifiable features of the swarm are indicated (see methods for definition). (**C**) Circularity plot for swarm expansion on LB, BHI, M8, M9, BM2 and PGM swarm agar (See Videos S1 to S6). End of swarm lag is indicated. Histogram for (**D**) swarm area, (**E**) perimeter, (**F**) radius of minimum bounding circle, (**G**) normalized area, and (**H**) number of growing tips. Pairwise comparison between every two media was performed by Tukey test. (P>0.05, ns; P<0.05, *; p<0.01, **, P<0.001 ***).

We utilized one-way ANOVA to isolate features which account for variance across swarm patterns observed on four media which promote dendritic swarming. All 10 macroscale features - Area (***), perimeter (***), RMBC (***), normalized area (***), branch length (**), branch angle (***), branch width (***), number of levels (*), number of primary branches (*), and number of growing tips (***) - could explain the variance across swarms on these media. Further, we utilized Tukey’s post hoc test to identify features that vary significantly between any two media (Figure 2D-H, Figure S1). For example, perimeter and area coverage (Mean ± SEM) values for swarm were significantly different (Figure 2D) for most pairwise comparisons, as well as the number of growing tips and normalized area. However, to understand the contribution of each feature to the swarm plasticity across different media, we performed a principal component analysis (PCA). For the PCA, we used all the ten macroscale features other than circularity of several swarms on each of the four media (methods). Principal component 1 (branch angle, area, perimeter, normalized area, and growing tips) contributed 37% to the variance, while component 2 (branch width and radius of minimum bounding circle) contributes to 25% (Figure 3A) of the variance across media. Principal Component 3 (branch length, number of primary branches and number of levels) contributed only 14% to the variance (Figure 3B). Normalized area versus perimeter could also distinguish PA14 swarm patterns on M8, M9, BM2, and PGM agar into distinct centroids (Figure 3C). We have used these two features in the rest of this study. Taken together, we have defined multiple quantifiable features of *P. aeruginosa* PA14 swarm that can be used to analyse perturbations to swarm patterns.

**Figure 3.**
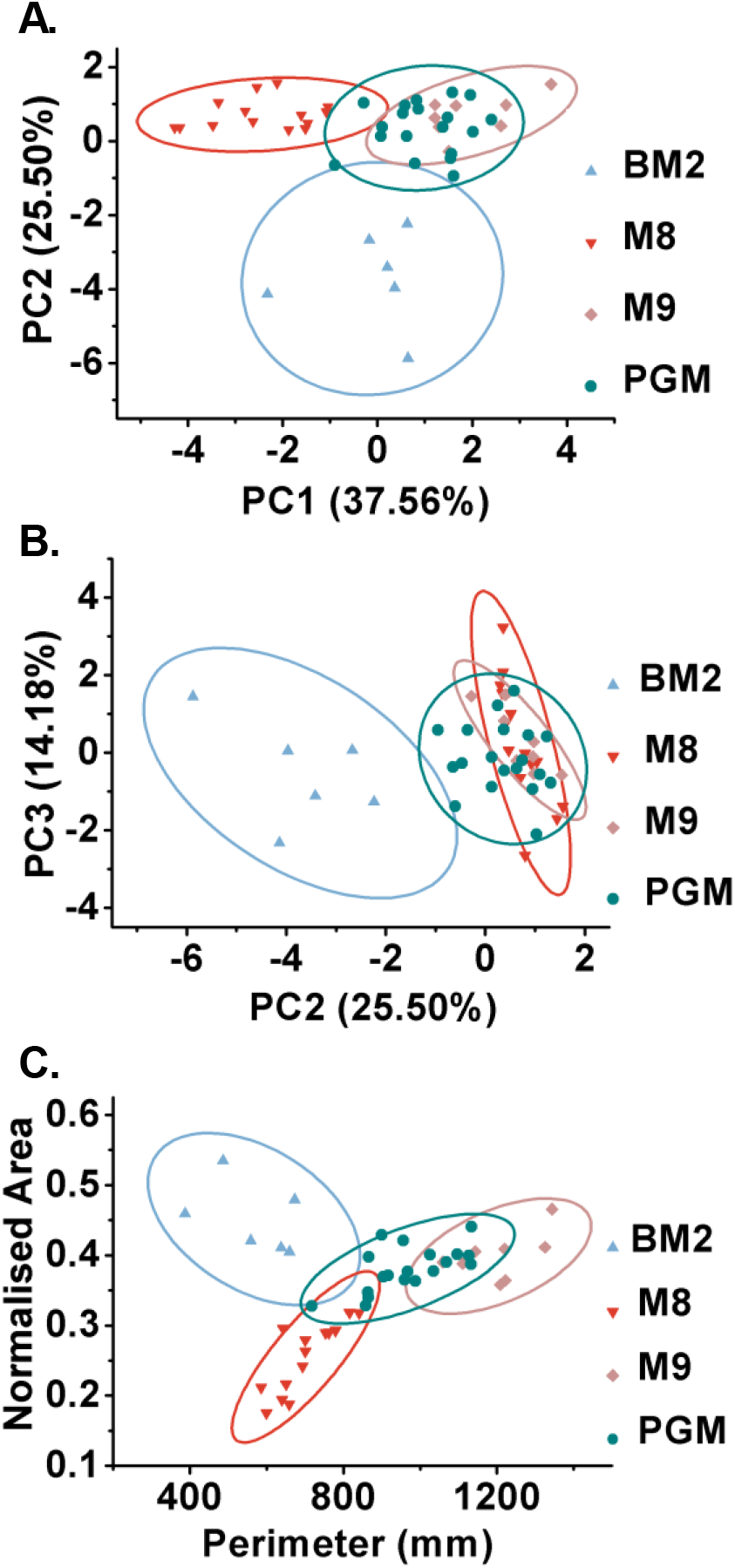
Principal component analysis of swarm features. (**A**) PC1 (branch angle, area, perimeter, normalized area and growing tips) and PC2 (branch width and radius of minimum bounding circle), and (**B**) PC2 vs. PC3 (branch length, number of primary branches and number of levels) plot for dendritic swarm pattern on BM2, M8, M9 and PGM agar. (**C**) Perimeter vs. normalized area plot for swarm patterns on BM2, M8, M9 and PGM agar. Centroids for each medium specific swarm are indicated in A, B and C.

### Several two-component genes of *P. aeruginosa* are conditional modulators of swarming

Media-dependent plasticity in PA14 swarming patterns strongly suggested that nutritional cues, in the growth media, might be critical for inducing swarming in *P. aeruginosa*. We then set out to ask how does *P. aeruginosa* sense such changes to execute swarming? In prokaryotes, the two-component systems (TCSs) are predominantly involved in sensing environmental signals such as nutrition, change in pH, redox state, osmolarity, light etc (Laub & Goulian, 2007; Stock, Robinson, & Goudreau, 2000; Zschiedrich, Keidel, & Szurmant, 2016). *P. aeruginosa* PA14 genome encodes 160 TCS genes (Lee et al., 2006; Liberati et al., 2006) and thought to provide exceptional adaptability of this bacterium to various environmental niches and a wide range of hosts including human, Drosophila, *C. elegans* and plants (Barreteau et al., 2009; Clatworthy et al., 2009; D’argenio et al., 2001; Francis et al., 2017; Rodrigue et al., 2000; Tan et al., 1999).

In a previous screen for genetic regulators of swarming in *P. aeruginosa* PA14, 12 TCS gene candidates were found (Yeung et al., 2009) to regulate swarming on BM2 agar. We hypothesized that two-component sensor kinases sense nutrient limitation and promote swarming. Since the media used in this study vary in both macro and micro-nutrients (Table S1), we expected to find different TCS genes to be required for swarming on different media. To test this hypothesis, we performed a targeted genetic screen for swarming using transposon insertion mutants for 113 PA14 TCS encoding genes (Liberati et al., 2006). The screen was performed on all 6 media, in duplicates. To assess the effect of TCS genes on swarming, we extracted three macroscale features - area, perimeter, and normalized area - from 681 swarms using MATLAB (see Figure S2 and S3). As shown in perimeter versus normalized area plot for four media in Figure 3A-D, many TCS genes were required for swarming (Table S2). We termed these <u>b</u>acterial <u>sw</u>arming (*bsw*) class of genes. We annotated unstudied sensor kinases as *bswS*, response regulators as *bswR*, and hybrid sensor kinases as *bswH* genes (Figure S4, and Table S3).

We found that four TCS genes were essential regulators of swarming. While nine TCS genes were required for swarming on at least two media. However, the largest number of TCS genes (31 out of 44) displayed medium specific role in swarming. Notably, 13 sensor kinases (SK) and 7 response regulators (RR) encoding genes were essential for swarming exclusively on BM2 agar, only 3 RRs and 2 SKs encoding genes were found essential for swarming on M8 agar alone. Swarming on PGM agar required 6 TCS genes while a single TCS gene encoding an RR was found essential exclusive to swarming on M9 agar. Media dependent requirement of TCS is displayed in a Venn diagram (Figure 4E). These observations suggest that swarming on the BM2 medium is dependent on signaling from many sensor kinases while swarming on other media is less reliant on them. It is also interesting to point out that in 23 cases either the response regulator or the sensor kinase, but not the both, was enriched in the screen – indicating a possible crosstalk in *P. aeruginosa* TCS signaling for swarming (Table S3). However, we did find 4 cognate pairs which can displayed phenotypic correlation for swarming.

**Figure 4.**
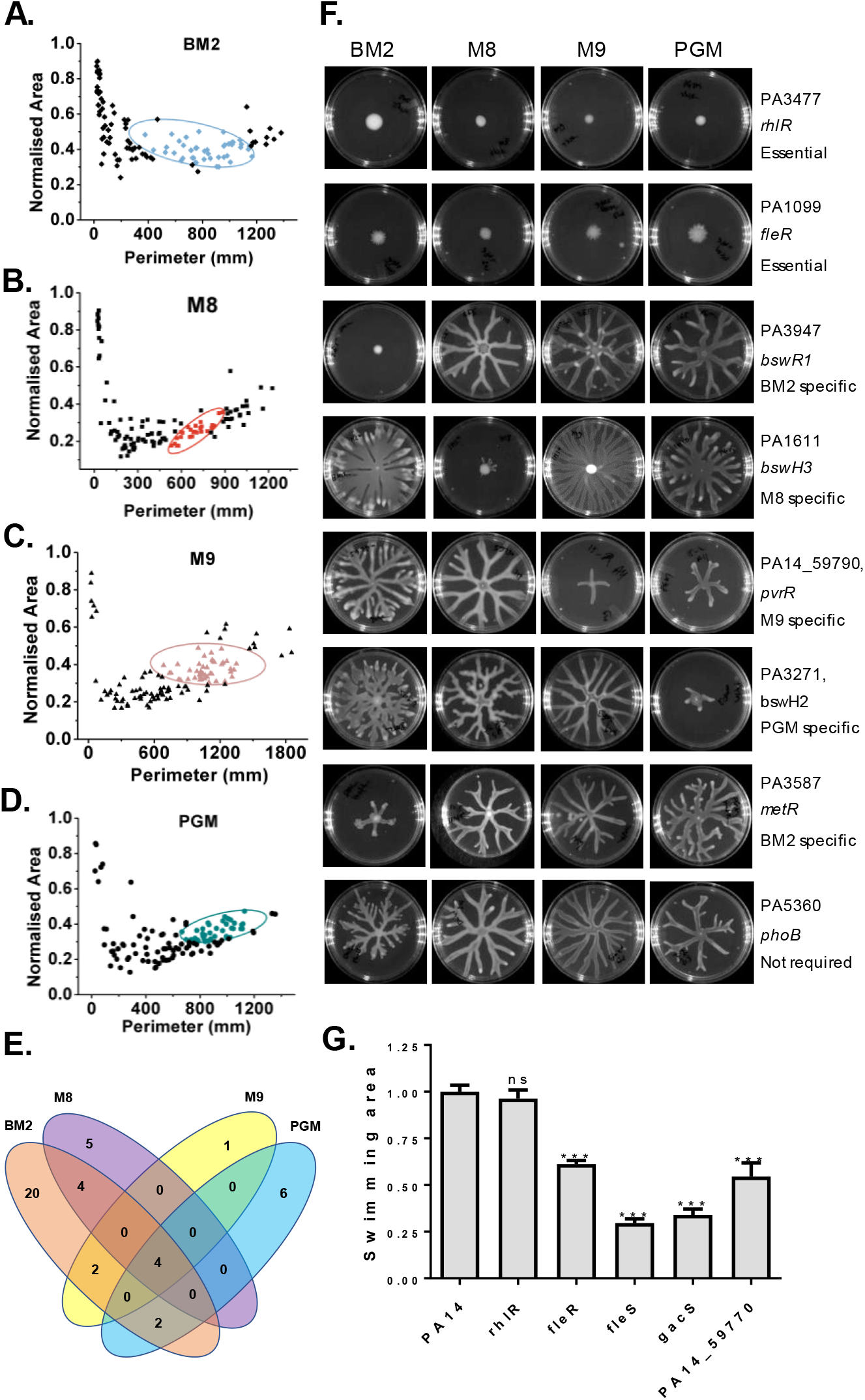
Two component systems of *P. aeruginosa* regulate swarming. Perimeter vs normalized area plot for PA14 and 113 TCS mutants on (**A**) BM2 agar, (**B**) M8 agar, (**C**) M9 agar, and (**D**) PGM agar. Centroid for wildtype PA14 on respective media is shown. All mutants outside the centroids represent weak swarmer or non swarmers. (**E**) Venn diagram to show media specific non-swarmer TCS mutants (also see Table S2). (**F**) Representative images of swarm of 6 TCS mutants, *rhlR* and *metR* strains on BM2, M8, M9 and PGM media. (**G**) Swimming phenotype of essential swarm regulators on LB-0.3% swim agar. Mean values were compared by unpaired *t* test (P>0.5, ns, p<0.001, ***).

Majority of TCS mutants displayed a strong, but context-dependant, swarming phenotype across media. A few striking examples are presented in Figure 4F. PA3947/rocR had a BM2 agar specific function (Figure 4F). Similarly, PA3271/*bswH2* and PA1611/*bswH3* had specific function in swarming on PGM agar and M8 agar respectively. PA14_59790/*pvrR*, a response regulator found exclusively in *P. aeruginosa* PA14 and PA7 genomes, showed a role in swarming on M9 and weakly on PGM agar (Figure 4F). As a control, we checked the context dependent requirement for PA3587/*metR*, a previously described transcriptional regulator of swarming on BM2 agar (Yeung et al., 2009). We found that *metR* was indeed swarming deficient on BM2 agar but swarming proficient on M8, M9 and PGM agar (Figure 3F). Interestingly, *phoB* required for swarming in low phosphate (2 mM) BM2 medium (Bains, Fernandez, and Hancock, 2012) was swarming proficient on all four media we tested (Figure 3F). This was expected as all media we used contain phosphate concentration of 24 mM or above.

Flagella and quorum sensing (QS) are essential for swarming (Kohler et al., 2000). Indeed, quorum defective mutant, *rhlR*, was a non-swarmer on all media we tested. However, *rhlR* had a wild-type swimming phenotype (Figure 4G). In contrast, essential swarming regulators of the TCS class were swimming defective (Figure 4G). *fleS* and *fleR* mutant showed moderate to severe swimming defect (Figure 4G), as also shown earlier (Ritchings et al, 1995). An analysis of transposon insertions in 26 flagellar genes (Liberati et al, 2006) also showed non-swarming phenotype (data not shown). Taken together, these experiments suggested that swimming ability is indeed essential for swarming under all conditions.

In all, our observations suggest that 44 TCS genes are required for swarming. This also indicates that many nutritional/ environmental cues promote swarming by activating specific two-component signalling circuits.

### Bsw genes differentially regulate swarming and biofilm formation

Biofilm and swarming constitute sessile and motile population, respectively; both of which rely on quorum sensing. In *P. aeruginosa* certain cellular components such as flagella are required both for swarming and for biofilm formation (Kohler et al., 2000; O’Toole & Kolter, 1998). In contrast, some regulatory components important for biofilm formation negatively regulate swarming motility (Akihiro Ueda and Thomas K. Wood, 2009; Bhuwan et al, 2012; Kuchma et al., 2007). For instance, higher cellular level of cyclic di-GMP molecule is considered a major switch for biofilm formation in many bacteria, including *P. aeruginosa* (Baker et al., 2016; Romling, Galperin, & Gomelsky, 2013; Valentini & Filloux, 2016) while suppressing swarming. Indeed, some of the genes discovered as swarm regulator in our study - gacA/gacS, *retS*, *sagS*, *bfiS*, *wspR* and *hptB* - are known to be regulators of biofilm formation (reviewed in Francis et al., 2017). This raised the question whether *bsw* genes discovered in this study regulate biofilm formation in PA14 and in what manner.

To understand the impact of *bsw* genes on biofilm formation, we assayed production of extracellular polymeric substances (EPS) by PA14 wild-type and *bsw* mutants as described (O’Toole, 2011). We used M63 medium recommended for quantification of biofilm in addition to M8, M9, BM2 and PGM broth (Table S4). Figure 5A shows swarm perimeter of all 44 swarm mutants (Table S2) and *rhlR* mutant on 4 different media in coloured disk format. Figure 5B shows crystal violet stain of *bsw* mutant on 5 media in coloured disk format. We found that eps production was not influenced by a change in media (compare disk colour in each column in 4B, values in Table S4). Only 3 mutants were unable to form the biofilm on all media (dark blue disks, Figure 4B). These included *fleS* and *fleR* mutants defective in flagella biogenesis, and *wspR*. However, four bsw mutants (*cheA, creC, cpxA, and tctE*) showed hyper-biofilm phenotype (deep yellow discs in Figure 4B), and thus appear to be negative regulators of biofilm formation, in a media independent manner. There were few media specific regulators of biofilm formation as well. For instance, *ntrB* was essential for EPS production in M63 and PGM broth while *kinB* was found essential only in M8 broth. The relationship between biofilm formation and swarm formation phenotypes is displayed in a double-faced Janus droplet map for four media in Figure 5C. The left half of the droplet represents swarm phenotype, while the right half represents swarming phenotype under the same condition (medium). We found that in 38 instances non-swarmer *bsw* mutants or a weak swarmer produced significantly higher amounts of eps (2-5 fold) than the wild-type PA14 (16 on BM2, 8 on PGM, 9 on M9, 5 on M8 and 8 on PGM). Taken together, the systematic analysis of swarm and EPS production on 4 different media (176 one on one comparisons) indicated that several TCS circuits regulate switch between swarming and biofilm formation.

**Figure 5.**
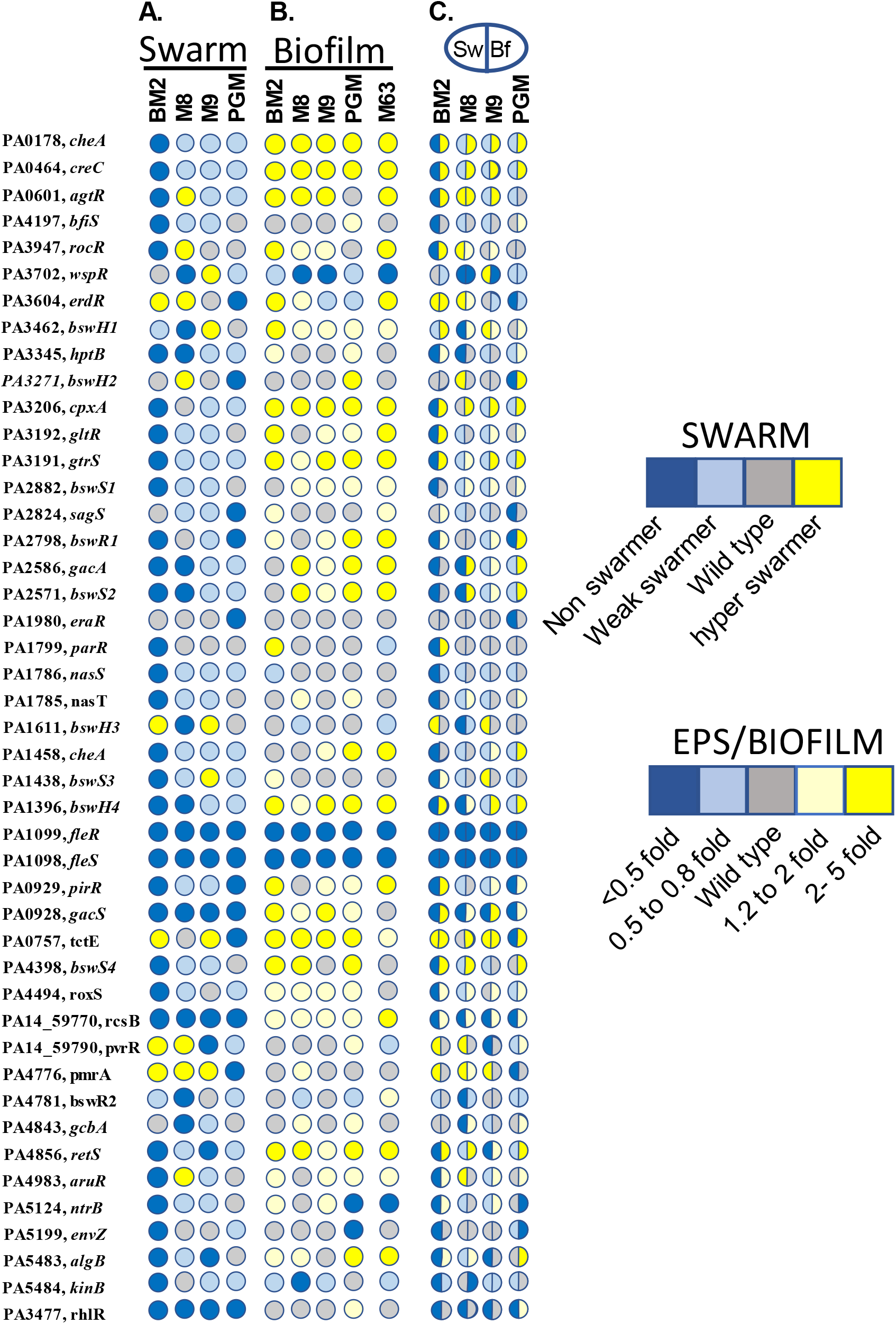
Media dependent role of TCS genes in swarming and biofilm formation. (**A**) Disk heat map of swarm phenotype of 44 TCS mutants and *rhlR*. Heat map is based on Perimeter values in Table S2. Swarm phenotype (non-swarmer, perimeter less than 20% of PA14 swarm perimeter; weak swarmer, >20% but less than mean - SD of PA14 perimeter; hyper swarmer, > mean + SD of PA14 swarm perimeter). (**B**) Extracellular polymeric substance (EPS) produced by 44 TCS mutants and *rhlR* in BM2, M8, M9, PGM and M63 broth, measured by crystal violet stain (values in Supplementary Table 4). (**C**) Janus droplet representation of media dependent biofilm and swarm phenotype. Left face shows swarming phenotype, while the right face reflects biofilm phenotype. Crystal violet staining for EPS quantification is represented as fraction of EPS production by PA14.

In summary, this study provides evidence that many TCS genes are critical for swarming in *P. aeruginosa* in a contextual manner (Figure 6). We find that PA14 swarming under one condition such as BM2 agar requires input from several TCS systems while swarming on M8, M9 and PGM media (condition II) rely on fewer TCS circuits. Nutritionally rich media, LB and BHI, do not support dendritic swarming in *P. aeruginosa* PA14. Thus, extrinsic nutritional cues in conjunction with bacterial SK/RR systems are critical in the modulation of *P. aeruginosa* swarming.

**Figure 6.**
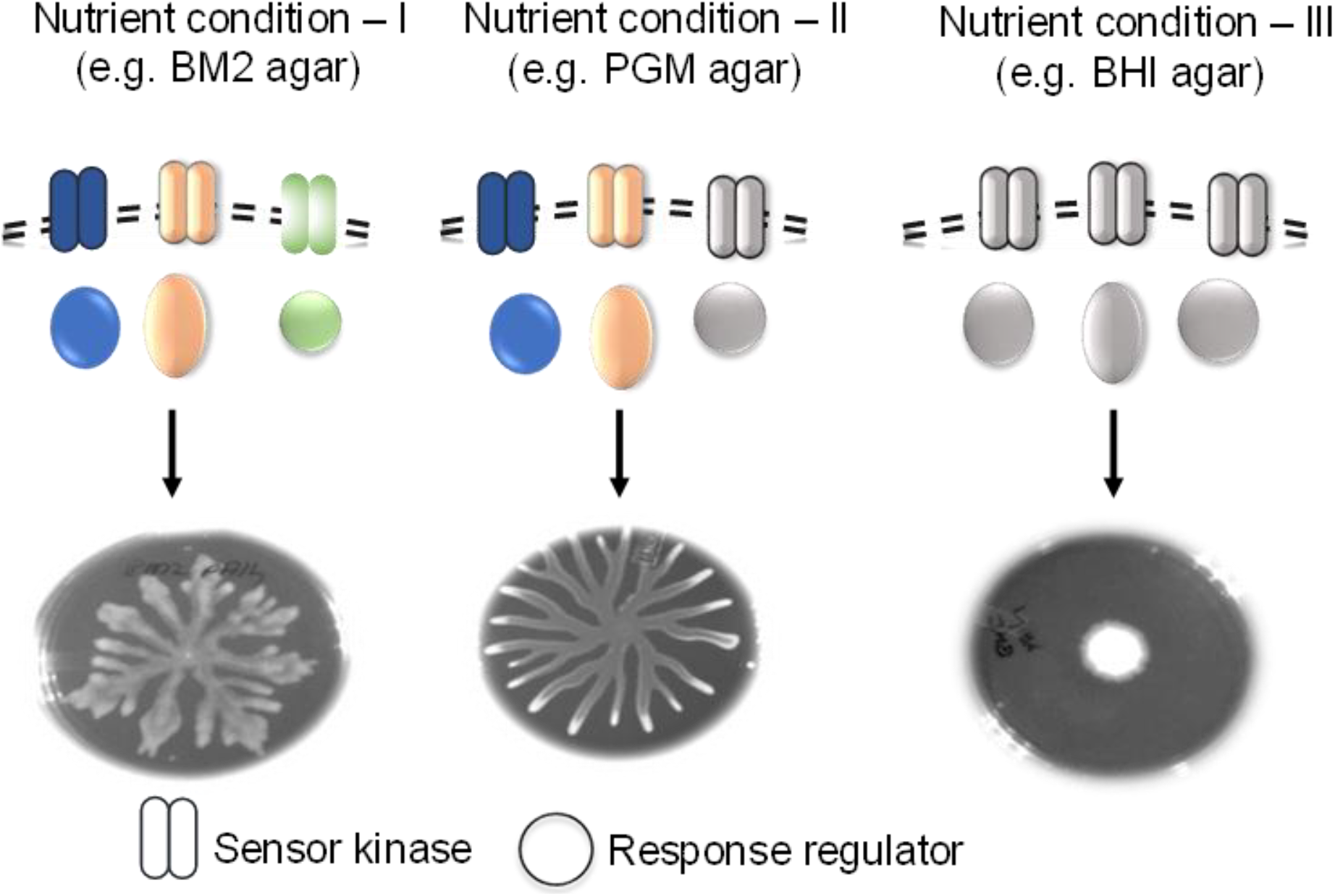
Media dependent plasticity in *P. aeruginosa* swarming. Growth media vary in macro and micronutrient contents. Some growth media (condition I) dependent on several TCS modules to allow swarming of *P. aeruginosa*. Another set of media (condition II) promote swarming but rely on a smaller but specific set of TCS genes for swarming. Certain other growth media (condition III) do not allow dendritic swarming of *P. aeruginosa.*

## DISCUSSION

In this study, we show that *P. aeruginosa* exhibits a remarkable, context-dependent plasticity in its swarming behaviour. This arises due to nutrient limitation in growth media and sensed by the TCS class of sensor kinases and their partners called response regulators. We provide a number of macroscale features of *P. aeruginosa* swarm to differentiate media specific swarm patterns into distinct populations.

We establish that swarm lag, the time to initiate branching or dendrite formation, is a quantifiable feature of *P. aeruginosa* swarming that can be represented as a change in circularity in time. To our surprise, swarm lag was significantly different between M8, M9, BM2 and PGM agar. The circularity remained close to 1 for the entire duration in non-dendritic growth on LB and BHI, again making it a valuable feature. Circularity, as well as other macroscale features, can be utilized to quantify the effect of the environmental factors or genetic regulators on swarm pattern.

One important question raised by this study is: What are the nutritional cues that promote swarming? Boyle et al (Boyle *et al*, 2015) have suggested that iron limitation is a requirement for swarming. Among the four media we used, only BM2 had the iron supplement. However, it does support PA14 swarming with a medium-specific pattern. The nitrogen content of the BM2 medium, however, is lower than that of M8 and M9 media (Table 1) suggesting, that nitrogen limitation could be a driver for initiating swarming under BM2 condition. Indeed, nitrogen related two-component systems - *ntrB* as well as *nasS*/*nasT* - were essential for swarming on BM2 media alone. These mutants also displayed weak swarming on other three media suggesting that nitrogen limitation may be a contributing factor to swarming in those media as well. Phosphate limitation is also a driver for swarming (Bains M, Fernández L, 2012) but it was not relevant for the four media we tested due to presence of phosphate in the media as well as dispensability of phosphate specific TCS, *phoB* and *phoR* (Table S1, Figure 4F) for swarming in this study. Involvement of low Mg++ and cationic peptides inducible *PmrA* on swarming on PGM agar suggests that Mg++ and cationic peptides may become relevant on certain media. Addition or removal of specific macro and micronutrients (Table S1) to/from these four media will serve to decipher additional nutritional cues that drive *bsw* TCS genes to modulate swarming. Do similar drivers exist in a *P. aeruginosa* infection setting in humans? A careful analysis of nutrients in body fluids of the host in different pathologies such as cystic fibrosis and diabetic foot ulcer can better out insight into *P. aeruginosa* pathology.

Our study raises a second question: How does nutrition impact swarming? It could be via modulation of flagellar output, modulation of rhamnolipid production or through novel pathways required for hitherto unidentified effector molecules for swarming. In planktonic growth, rhamnolipid production is induced at the end of the lag phase which corresponds with nutrient limitation (Caiazza, Shanks, & O’Toole, 2005). In addition, Xavier *et al* have shown that rhamnolipid production is regulated by nitrogen limitation in minimal media (Xavier et al., 2011). We find that rhamnosyltransferase chain A (RhlA) transcription in much higher in PGM broth than in LB broth. Transcription of *rhlA* is further induced on PGM-0.6% swarm agar than PGM broth (data not shown). This indicated that rhamnolipid production is dependent on media as well as surface contact. Careful analyses of rhamnolipid production in response to removal or limitation of specific nutrient one at a time would be instructive in understanding the regulation of swarming. Some of the TCS components may regulate swarming via the modulation of cyclic di-GMP levels. We found that PA4398 is a positive regulator of swarming on BM2 medium and a negative regulator of EPS production (Figure 5A-B). This is agreement with an earlier report where the mutation in PA4398/bswS4 led to a 50% increase in intracellular cyclic-di-GMP which is linked to enhanced EPS production (Strehmel et al., 2015). Indeed, six response regulators (*RocR, WspR, ErdR, PvrR*, PA4781, and PA4843) we identified as *bsw* genes do possesses diguanylate cyclase (GGDEF) or phosphodiesterase (EAL and HD-GYP) domains for regulation of c-di-GMP turnover in bacteria. There also appears to be a potential for the role of nitrogen limitation in swarming. A TCS pair (NasS/nasT) involved in nitrate assimilation is essential for swarming on BM2. In addition, sensor kinase NtrB and two ntrC domain containing response regulator are also essential for swarming (Table S3). A systematic study of each TCS component in a medium-specific context will help decipher the possible mechanism of modulation of swarming. Over time, it would be possible to build up the regulatory program for swarming in *P. aeruginosa*.

We were surprised to find 32 TCS modulators of swarming on BM2 agar in comparison to 12 genes uncovered in a previous screen (Yeung et al., 2009). We believe that additional TCS regulators of swarming were found due to the utilisation of a large (90 mm) dish format for each strain in our study in comparison to the 96 well multiplicator format deployed for the primary screen in the previous report (Yeung 2009). We find that avoidance zone between two swarms occurs at about 5 mm distance and 96 prong multiplicator does not allow enough expansion of swarm to detect all non-swarmers in our laboratory (data not shown). A recent study from our laboratory shows that *P. aeruginosa* can detect both bacteria and non-biological obstacles on PGM swarm agar (Harshitha et al., 2018) reiterating that swarming in *P. aeruginosa* is sensitive to environmental cues including proximity to isogenic bacteria.

*P. aeruginosa* genome exhibits expansion of the two-component system encoding genes to 160 one of the largest repertoires among eubacteria. The GacS network, along with HptB and SagS branches control biofilm, virulence, and motility. We found that six components of this extended network (*gacS*, *gacA*, *retS*, PA1611/*bswH3*, *hptB*, *sagS* and *bfiS*) were essential or conditional regulators of swarming in *P. aeruginosa*. On the other hand, some of the conditional swarm modulators uncovered in this study are linked to nutrient assimilation (eg *ntrB*, *nasS* and *nasT*). Several TCS mutants previously described as motility, swarming or biofilm-related loci including *PA14_16500* (*wspR*) (Chen et al., 2014), *PA14_20800* (*hptB*) (Bhuwan et al., 2012; Hsu, Chen, Peng, & Chang, 2008) *PA14_59770* (*rcsB*) (Giraud, Filloux, & Mikkelsen, 2009) *PA14_59790* (*pvrR*) (Giraud et al., 2009; Zheng et al., 2016) *PA14_57170* (Strehmel et al., 2015) enriched as swarm regulator, at least under one condition, in our study. To the best of our knowledge, the others are not to have any known swarming-related function. Based on our study, we would like to propose that some of the *P. aeruginosa* TCS genes may have evolved, and been retained in the genome, to modulate swarming motility.

One of the surprises from this study came in the form of antagonistic regulation of swarm and biofilm formation by TCS genes. There were 38 cases of an inverse relationship between biofilm and swarm. This indicates that several TCS signaling circuits might promote swarming while suppressing biofilm formation. This leads to an interesting hypothesis that *P. aeruginosa* can possibly only exist in one state (either swarming or biofilm) at one time. Swarming bacteria are believed to be antibiotic resistant (Butler, Wang, & Harshey, 2010; Overhage et al., 2008). One of the directions to pursue will be to ask if swarming population of *P. aeruginosa* is more susceptible to antibiotics than *P. aeruginosa* in a biofilm. If so, could we change the state of the bacteria and make them more susceptible to antibiotics. This could be done by pharmacological intervention or simply by perturbation of host body fluids? A comprehensive analysis of transcriptional events that initiate biofilm versus initiate swarm will add to better understanding of the differences between these two quorum-dependent processes.

## MATERIALS AND METHODS

### Bacterial strains and growth conditions

*Pseudomonas aeruginosa* PA14 was used as the wild-type strain. All the mutant strains used in this work are transposon insertion library of PA14 (Liberati et al., 2006). Unless otherwise mentioned, all strains were grown in Luria Bertani (LB) broth under standard laboratory conditions. For selection of transposon mutants, gentamycin (50 μg mL^−1^) antibiotic was used. No antibiotics were used in the swarm dishes. Additional growth media used in this study are BHI, BM2, M9, M8 (a modified M9 media without NH_4_Cl and CaCl_2_), PGM and M63 medium (100 μM KH_2_PO_4_, 15.14 mM (NH_4_)_2_SO_4_, 0.36 μM FeSO_4._H_2_O, 1 mM MgSO_4_ and 4% arginine). Media composition for the rest is provided in Table 1.

### Swarming assay and screening

Swarming motility assays were performed as previously described (Overhage et al., 2008; Yeung et al., 2009), with additional modifications. Appropriate medium (LB, BHI, M8, M9, BM2 or PGM) was solidified with 0.6% Bacto^™^ agar (BD) and inoculated after 16 hours. All plates were inoculated at the centre with 2 μL of overnight bacterial culture in LB broth (OD_600_ = 2.8 - 3.0) and incubated at 37°C for 24 hours. All the no-swarmer phenotypes were confirmed at least in three independent experiments.

### Swimming Assay

For swimming tests, PGM containing 0.3% Bacto^™^ agar (BD) was used. A 5 μl of 2 ml overnight culture was inoculated into 5 ml LB (secondary culture) and incubated at 37°C for 5 hours (or OD_600_ = 1). Using sterile toothpicks, the secondary culture was introduced into the centre of swim agar plate by puncturing into the agar but without touching the base of the plates. Plates were incubated at 37°C for 24 hours right side up. Swimming proficiency were analysed by measuring the swim area covered.

### Biofilm quantification

Biofilm formations were assayed as previously described (O’Toole, 2011). Briefly, *P. aeruginosa* strains were grown overnight in 2 mL of LB broth at 37°C. Overnight culture was then diluted with biofilm media (M63) or swarming media (M9, M8, BM2 or PGM) at ratio of 1:100. 100 μL of the dilution were added to each well of a 96-well microtiter plate, in triplicate. Microtiter plate was then incubated at 37°C for 24 hours. After incubation, the plates were rinsed with tap water to remove the planktonic cells and air dried. 0.1% solution of crystal violet in water was added to each well of microtiter dish for staining, followed by 15 minutes incubation at room temperature. After rinsing the excess stain by vigorous shaking and tapping of the plate on paper towels, plates were dried for a few hours. Crystal violent stain in each well was solubilized in 125 μL 30% acetic acid followed by 15 minutes incubation at room temperature. Biofilm formation was quantified by measuring absorbance at 550 nm. PA14 WT was taken as reference; a mutant strain for EPS, *pelA*, was taken as control. The results are represented in table S4 as mean value (standard error of mean) and sample size (n). All strains were stained 3 - 4 independent times, with three replicates on every occasion.

### Principal component analysis for swarm features

Principal component analysis (PCA) has been performed on different parameters of the colony pattern of wild type PA14 in different nutrient media. We divide these patterns into two classes branching and non-branching patterns. Wild type PA14 produces non-branching patterns on LB and BHI swarm agar but produces branching patterns on BM2, M8, M9 and PGM swarm agar. We defined 10 parameters to describe a branching pattern as follows:

Branch length (BL): Length of all dendrites in the pattern
Branch Angle (BA): Angle between two dendrites in the pattern
Branch width (BW): Dendrite width of all branches in the pattern
Area (A): Area covered by the swarm
Perimeter (P): Perimeter of the pattern formed by the swarm
Radius of Minimum bounding circle radius (RMBC): Radius of the circle that can encircle the pattern
Number of levels (NL): Number of levels of branching a dendrite undergoes
Normalised Area (NA): Ratio of Area of the pattern and Area of the minimum bounding circle
Number of primary branches (NPB): Number of branches originating from the point of inoculation
Growing tips (GT): Total number of dendrite tips present in the swarm at 24 hours.

The non-dendritic patterns (on LB and BHI agar) are characterized by Area (A), Perimeter (P), Radius of minimum bounding circle (RMBC), and the Normalized Area (NA). We chose two independent parameters to differentiate patterns of dendritic swarm. The parameters chosen i.e. Perimeter and Normalised area are also defined for patterns on LB and BHI.

### Statistical analysis

One-way ANOVA was used to analyse variance of swarming traits across swarms on different media. For comparison of means of traits, post hock Tukey test was performed in Figure 2. In all other cases, unpaired student’s t test was performed to compare mean ± SEM as indicated.

#### Supplementary files

Tables (S1 to S4)

Figures (S1 to S4)

Videos (S1 to S6)

## ACKNOLEDGMENT

*Pseudomonas aeruginosa* PA14 transposon insertion library was a gift from Prof. Frederick M. Ausubel, Massachusetts General Hospital, Boston. This work was supported by the Wellcome Trust/DBT India Alliance Fellowship (Grant no. IA/I/13/1/500919) awarded to Varsha Singh and Robert Bosch Innovation Centre Grant (Grant no. RBCO0014) awarded to Manoj Varma and Varsha Singh. We thank Deepak K. Saini, Sandhya S. Visveswaraiah and Sambuddho Mukherjee for critical reading of the manuscript.

## STATEMENT OF COMPETING INTERESTS

Authors declare no conflict of interest.

## REFERENCES

Akihiro Ueda and Thomas K. Wood. (2009). Connecting Quorum Sensing, c-di-GMP, Pel Polysaccharide, and Biofilm Formation in Pseudomonas aeruginosa through Tyrosine Phosphatase TpbA, 5(6), 1–15. http://doi.org/10.1371/journal.ppat.1000483

Bains M, Fernández L, H. R. (2012). Phosphate Starvation Promotes Swarming Motility and Cytotoxicity of Pseudomonas aeruginosa, 78(18), 6762–6768. http://doi.org/10.1128/AEM.01015-12

Baker, A. E., Diepold, A., Kuchma, S. L., Scott, J. E., Ha, D. G., Orazi, G., … O’Toole, G. A. (2016). A PilZ domain protein FlgZ mediates c-di-GMP-dependent swarming motility control in *Pseudomonas aeruginosa*. Journal of Bacteriology, 198(April), JB.00196-16. http://doi.org/10.1128/JB.00196-16

Barreteau, H., Bouhss, A., Fourgeaud, M., Mainardi, J. L., Touzé, T., Gérard, F., … Mengin-Lecreulx, D. (2009). Human- and plant-pathogenic Pseudomonas species produce bacteriocins exhibiting colicin M-like hydrolase activity towards peptidoglycan precursors. Journal of Bacteriology, 191(11), 3657–3664. http://doi.org/10.1128/JB.01824-08

Bhuwan, M., Lee, H., Peng, H., & Chang, H. (2012). Histidine-containing Phosphotransfer Protein-B (HptB) Regulates Swarming Motility through Partner-switching System in Pseudomonas aeruginosa PAO1 Strain □, 287(3), 1903–1914. http://doi.org/10.1074/jbc.M111.256586

Boyle, K. E., Monaco, H., van Ditmarsch, D., Deforet, M., & Xavier, J. B. (2015). Integration of Metabolic and Quorum Sensing Signals Governing the Decision to Cooperate in a Bacterial Social Trait. PLoS Computational Biology, 11(6), 1–26. http://doi.org/10.1371/journal.pcbi.1004279

Butler, M. T., Wang, Q., & Harshey, R. M. (2010). Cell density and mobility protect swarming bacteria against antibiotics. Proceedings of the National Academy of Sciences, 107(8), 3776–3781. http://doi.org/10.1073/pnas.0910934107

Caiazza, N. C., Shanks, R. M. Q., & O’Toole, G. A. (2005). Rhamnolipids modulate swarming motility patterns of Pseudomonas aeruginosa. Journal of Bacteriology, 187(21), 7351–7361. http://doi.org/10.1128/JB.187.21.7351-7361.2005

Caiazza, N. C., Shanks, R. M. Q., & Toole, G. A. O. (2005). Rhamnolipids Modulate Swarming Motility Patterns of Pseudomonas aeruginosa, 187(21), 7351–7361. http://doi.org/10.1128/JB.187.21.7351

Chen, A. I., Dolben, E. F., Okegbe, C., Harty, C. E., Golub, Y., Thao, S., … Hogan, D. A. (2014). Candida albicans Ethanol Stimulates Pseudomonas aeruginosa WspR-Controlled Biofilm Formation as Part of a Cyclic Relationship Involving Phenazines, 10(10). http://doi.org/10.1371/journal.ppat.1004480

Clatworthy, A. E., Lee, J. S. W., Leibman, M., Kostun, Z., Davidson, A. J., & Hung, D. T. (2009). Pseudomonas aeruginosa infection of zebrafish involves both host and pathogen determinants. Infection and Immunity, 77(4), 1293–1303. http://doi.org/10.1128/IAI.01181-08

D’argenio, D. a., Gallagher, L. a, Berg, C. a, Manoil, C., & Argenio, D. a D. (2001). Drosophila as a model host for Pseudomonas aeruginosa infection. Journal of Bacteriology, 183(4), 1466–1471. http://doi.org/10.1128/JB.183.4.1466

Francis, V. I., Stevenson, E. C., & Porter, S. L. (2017). Two-component systems required for virulence in Pseudomonas aeruginosa. FEMS Microbiology Letters, 364(11), 1–22. http://doi.org/10.1093/femsle/fnx104

Giraud, C., Filloux, A., & Mikkelsen, H. (2009). Expression of Pseudomonas aeruginosa CupD Fimbrial Genes Is Antagonistically Controlled by RcsB and the EAL-Containing PvrR Response Regulators, 4(6). http://doi.org/10.1371/journal.pone.0006018

Harshey, R. M., & Matsuyama, T. (1994). Dimorphic transition in Escherichia coli and Salmonella typhimurium: Surface-induced differentiation into hyperflagellate swarmer cells. Proceedings of the National Academy of Sciences of the United States of America, 91(18), 8631–8635. http://doi.org/10.1073/pnas.91.18.8631

Harshitha S. Kotian, Shalini Harkar, Shubham Joge, Ayushi Mishra, Amith Zafal, Varsha Singh, M. M. V. (2018). Spatial Awareness of a Bacterial Swarm. BioRxiv. https://doi.org/10.1101/341529

Henrichsen, J. (1972). Bacterial surface translocation: a survey and a classification. Bacteriological Reviews, 36(4), 478–503.

Hsu, J., Chen, H., Peng, H., & Chang, H. (2008). Characterization of the Histidine-containing Phosphotransfer Protein B-mediated Multistep Phosphorelay System in, 283(15), 9933–9944. http://doi.org/10.1074/jbc.M708836200

Kaiser, D., & Warrick, H. (2011). Myxococcus xanthus swarms are driven by growth and regulated by a pacemaker. Journal of Bacteriology, 193(21), 5898–5904. http://doi.org/10.1128/JB.00168-11

Kearns, D. B. (2010). A field guide to bacterial swarming motility. Nature Reviews. Microbiology, 8(9), 634–44. http://doi.org/10.1038/nrmicro2405

Kearns, D. B., & Losick, R. (2003). Swarming motility in undomesticated Bacillus subtilis. Molecular Microbiology, 49(3), 581–590. http://doi.org/10.1046/j.1365-2958.2003.03584.x

Kohler, T., Curty, L. K., Barja, F., Van Delden, C., & Pechere, J. C. (2000). Swarming of Pseudomonas aeruginosa is dependent on cell-to-cell signaling and requires flagella and pili. Journal of Bacteriology, 182(21), 5990–5996. http://doi.org/10.1128/JB.182.21.5990-5996.2000

Kuchma, S. L., Brothers, K. M., Merritt, J. H., Liberati, N. T., Ausubel, F. M., & O’Toole, G. A. (2007). BifA, a cyclic-di-GMP phosphodiesterase, inversely regulates biofilm formation and swarming motility by Pseudomonas aeruginosa PA14. Journal of Bacteriology, 189(22), 8165–8178. http://doi.org/10.1128/JB.00586-07

Laub, M. T., & Goulian, M. (2007). Specificity in Two-Component Signal Transduction Pathways. Annual Review of Genetics, 41(1), 121–145. http://doi.org/10.1146/annurev.genet.41.042007.170548

Lee, D. G., Urbach, J. M., Wu, G., Liberati, N. T., Feinbaum, R. L., Miyata, S., … Ausubel, F. M. (2006). Genomic analysis reveals that Pseudomonas aeruginosa virulence is combinatorial. Genome Biology, 7(10). http://doi.org/10.1186/gb-2006-7-10-r90

Liberati, N. T., Urbach, J. M., Miyata, S., Lee, D. G., Drenkard, E., Wu, G., … Ausubel, F. M. (2006). An ordered, nonredundant library of Pseudomonas aeruginosa strain PA14 transposon insertion mutants. Proceedings of the National Academy of Sciences of the United States of America, 103(8), 2833–8. http://doi.org/10.1073/pnas.0511100103

Morales-Soto, N., Anyan, M. E., Mattingly, A. E., Madukoma, C. S., Harvey, C. W., Mark, A., … Shrout, J. D. (2015). Preparation, Imaging, and Quantification of Bacterial Surface Motility Assays. Journal of Visualized Experiments, (98), 1–10. http://doi.org/10.3791/52338

Morris, J. D., Hewitt, J. L., Wolfe, L. G., Kamatkar, N. G., Chapman, S. M., Diener, J. M., … Shrout, J. D. (2011a). Imaging and analysis of Pseudomonas aeruginosa swarming and rhamnolipid production. Applied and Environmental Microbiology, 77(23), 8310–8317. http://doi.org/10.1128/AEM.06644-11

Morris, J. D., Hewitt, J. L., Wolfe, L. G., Kamatkar, N. G., Chapman, S. M., Diener, J. M., … Shrout, J. D. (2011b). Imaging and Analysis of Pseudomonas aeruginosa Swarming and Rhamnolipid Production □ †, 77(23), 8310–8317. http://doi.org/10.1128/AEM.06644-11

O’Toole, G. A. (2011). Microtiter Dish Biofilm Formation Assay. Journal of Visualized Experiments, (47), 10–11. http://doi.org/10.3791/2437

O’Toole, G. A., & Kolter, R. (1998). Flagellar and twitching motility are necessary for Pseudomonas aeruginosa biofilm development. Molecular Microbiology, 30(2), 295–304. http://doi.org/10.1046/j.1365-2958.1998.01062.x

Overhage, J., Bains, M., Brazas, M. D., & Hancock, R. E. W. (2008). Swarming of Pseudomonas aeruginosa is a complex adaptation leading to increased production of virulence factors and antibiotic resistance. Journal of Bacteriology, 190(8), 2671–2679. http://doi.org/10.1128/JB.01659-07

Overhage, J., Lewenza, S., Marr, A. K., & Hancock, R. E. W. (2007). Identification of Genes Involved in Swarming Motility Using a Mutant Library, 189(5), 2164–2169. http://doi.org/10.1128/JB.01623-06

Patrick, J. E., & Kearns, D. B. (2009). Laboratory strains of Bacillus subtilis do not exhibit swarming motility. Journal of Bacteriology, 191(22), 7129–7133. http://doi.org/10.1128/JB.00905-09

Rashid, M. H., & Kornberg, A. (2000). Inorganic polyphosphate is needed for swimming, swarming, and twitching motilities of Pseudomonas aeruginosa, 97(9), 4885–4890.

Ritchings, B. W., Almira, E. C., Lory, S., & Ramphal, R. (1995). Cloning and phenotypic characterization of fleS and fleR, new response regulators of Pseudomonas aeruginosa which regulate motility and adhesion to mucin. Infection and Immunity, 63(12), 4868–4876.

Rodrigue, A., Quentin, Y., Lazdunski, A., Méjean, V., & Foglino, M. (2000). Two-component systems in Pseudomonas aeruginosa: why so many? Trends in Microbiology, 8(11), 498–504. http://doi.org/10.1016/S0966-842X(00)01833-3

Romling, U., Galperin, M. Y., & Gomelsky, M. (2013). Cyclic di-GMP: the First 25 Years of a Universal Bacterial Second Messenger. Microbiology and Molecular Biology Reviews, 77(1), 1–52. http://doi.org/10.1128/MMBR.00043-12

Shrout, J. D., Chopp, D. L., Just, C. L., Hentzer, M., Givskov, M., & Parsek, M. R. (2006). The impact of quorum sensing and swarming motility on Pseudomonas aeruginosa biofilm formation is nutritionally conditional, 62(October), 1264–1277. http://doi.org/10.1111/j.1365-2958.2006.05421.x

Singh, V., & Aballay, A. (2006). Heat-shock transcription factor (HSF)-1 pathway required for Caenorhabditis elegans immunity. Proceedings of the National Academy of Sciences of the United States of America, 103(35), 13092–7. http://doi.org/10.1073/pnas.0604050103

Stock, A. M., Robinson, V. L., & Goudreau, P. N. (2000). Two-component signal transduction. Annu. Rev. Biochem, 69, 183–215. http://doi.org/10.1146/annurev.biochem.69.1.183

Strehmel, J., Neidig, A., Nusser, M., Geffers, R., Brenner-Weiss, G., & Overhage, J. (2015). Sensor kinase PA4398 modulates swarming motility and biofilm formation in Pseudomonas aeruginosa PA14. Applied and Environmental Microbiology, 81(4), 1274–1285. http://doi.org/10.1128/AEM.02832-14

Sun, J., Singh, V., Kajino-Sakamoto, R., & Aballay, A. (2011). Neuronal GPCR Controls Innate Immunity by Regulating Non-Canonical Unfolded Protein Response Genes. Science, 332(6030), 729–732. http://doi.org/10.1126/science.1203411.Neuronal

Tan, M.-W., Mahajan-Miklos, S., & Ausubel, F. M. (1999). Killing of Caenorhabditis elegans by Pseudomonas aeruginosa used to model mammalian bacterial pathogenesis. Proceedings of the National Academy of Sciences, 96(2), 715–720. http://doi.org/10.1073/pnas.96.2.715

Tremblay, J., & Déziel, E. (2008). Improving the reproducibility of Pseudomonas aeruginosa swarming motility assays. Journal of Basic Microbiology, 48(6), 509–515. http://doi.org/10.1002/jobm.200800030

Tremblay, J., & Déziel, E. (2010). Gene expression in Pseudomonas aeruginosa swarming motility. BMC Genomics, 11(1), 587. http://doi.org/10.1186/1471-2164-11-587

Valentini, M., & Filloux, A. (2016). Biofilms and Cyclic di-GMP (c-di-GMP) signaling: Lessons from Pseudomonas aeruginosa and other bacteria. Journal of Biological Chemistry, 291(24), 12547–12555. http://doi.org/10.1074/jbc.R115.711507

Xavier, J. B., Kim, W., & Foster, K. R. (2011). A molecular mechanism that stabilizes cooperative secretions in Pseudomonas aeruginosa. Molecular Microbiology, 79(1), 166–179. http://doi.org/10.1111/j.1365-2958.2010.07436.x

Yeung, A. T. Y., Torfs, E. C. W., Jamshidi, F., Bains, M., Wiegand, I., Hancock, R. E. W., & Overhage, J. (2009). Swarming of Pseudomonas aeruginosa is controlled by a broad spectrum of transcriptional regulators, including MetR. Journal of Bacteriology, 191(18), 5592–5602. http://doi.org/10.1128/JB.00157-09

Zheng, Y., Tsuji, G., Opoku-temeng, C., & Sintim, H. O. (2016). Inhibition of P. aeruginosa c-di-GMP phosphodiesterase RocR and swarming motility by a benzoisothiazolinone derivative, 6238–6244. http://doi.org/10.1039/c6sc02103d

Zschiedrich, C. P., Keidel, V., & Szurmant, H. (2016). Molecular Mechanisms of Two-Component Signal Transduction. Journal of Molecular Biology, 428(19), 3752–3775. http://doi.org/10.1016/j.jmb.2016.08.003

